# DPAC: Prediction and Design of Protein-DNA Interactions via Sequence-Based Contrastive Learning

**DOI:** 10.1101/2025.05.14.654102

**Authors:** Leo Tianlai Chen, Rishab Pulugurta, Pranay Vure, Pranam Chatterjee

## Abstract

Interactions between DNA and proteins are pivotal in natural biological processes, and designing proteins that can bind to DNA with high specificity is crucial for advancing genomic technologies. Existing state-of-the-art models for both modeling and designing protein-DNA interactions primarily rely on structural information, facing limitations in scalability and efficiency for large-scale applications. Notable methods like AlphaFold 3 and RosettaTTAFold All-Atom exist, but they are inefficient and inherently struggle at modeling conformationally unstable proteins, such as transcription factors, which arguably represent the most important class of DNA-binding proteins. Here, we present **DPAC**^1^ (**D**NA-**P**rotein binding **A**lignment via **C**ontrastive learning), which leverages pre-trained protein and DNA language models via a contrastive loss to align the two modalities in a high-dimensional shared latent space. DPAC not only significantly accelerates the design process compared to current structure-based methods but also demonstrates a strong ability to differentiate real binders from non-binders. Our model achieves an AUC score of 0.591 on a low identity set, outperforming state-of-the-art structure-based methods. Additionally, DPAC integrates simulated annealing for the design of new protein sequences with optimized DNA binding affinity, successfully recovering binding affinity in engineered sequences by up to 20% in *in silico* tests. Our results highlight DPAC’s potential for facilitating the design and discovery of sequence-specific DNA-binding proteins, paving the way for advancements in genomic research and biotechnology applications.

## 1 Introduction

DNA binding proteins play essential roles in regulating gene expression, repairing DNA, and influencing critical biomolecular interactions (1). Recently, novel deep learning-based approaches to design and model DNA-binding proteins have emerged, but heavily rely on structural information to predict interactions and identify potential contact sites. These models, including RoseTTAFoldNA (RFNA) (2), followed by more general, atom-based models RoseTTAFold All-Atom (RFAA) (3) and AlphaFold 3 (4), have significantly accelerated molecular docking and design of DNA-binding proteins beyond expensive molecular dynamics simulations (5). However, these structure-based methods still face limitations in scalability and computational efficiency, particularly for large-scale virtual screening efforts. Additionally, they are limited to protein-DNA complexes that adopt stable conformations, precluding accurate modeling of large classes of DNA binding proteins, most specifically transcription factors, which adopt largely disordered conformations (6). Overall, the inherent complexity and variability in protein-DNA interactions demand robust models capable of accurately predicting binding specificity and affinity solely based on sequence information.

Recent advancements in deep learning and natural language processing have opened new avenues for protein and DNA sequence analysis. Pre-trained protein language models (pLMs), and to a lesser extent, nucleic acid language models, have shown success in capturing key physicochemical, structural, and functional features of input molecules (7; 8; 9; 10). However, leveraging these models to predict and design protein-DNA interactions remains a unsolved task, and no purely sequence-based protein-DNA design module has been developed.

To address this shortcoming, in this paper, we introduce **DPAC** (**D**NA-**P**rotein Binding **A**lignment via **C**ontrastive Learning), a new sequence-based approach that addresses these challenges. Our model utilizes state-of-the-art, pre-trained protein and DNA language models to create high-dimensional, feature-rich embeddings of sequences, which are then aligned using a CLIP-like two tower-based contrastive learning framework (11). DPAC further leverages fast simulated annealing for protein sequence generation, providing a principled method of searching the contrastive latent space to identify sequences conferring optimized DNA-specific binding properties. As such, DPAC not only predicts binding interactions but also facilitates the design of new optimized, DNA-binding protein sequences, making it a powerful tool for high-throughput virtual screening and protein design, and motivating downstream experimental utilization.

## 2. Methods

Protein-nucleic acid binding is the interaction between a protein sequence *o* and a DNA sequence *a*. A key question in biological research is how to identify which protein binds to a given DNA sequence *a* or engineer a protein for improved binding and vice versa. To address this, screening, whether virtual or experimental, is essential. Screening involves evaluating a set of protein sequences *{o*1, *o*2, … *}, o*_*n*_ and DNA sequences *{a*1, *a*2, …, *a*_*n*_*}* with a measurement score *S*(·). The goal is to maximize the score for real binding pairs and minimize it for decoy pairs. This problem is well-suited for contrastive learning, which aims to distance negative samples while keeping positive samples close in a high-dimensional space. Our model uses pre-trained language models and contrastive learning to better identify binding pairs and to virtually engineer protein sequences using simulated annealing.

### DPAC

Our model is fundamentally a two-tower architecture similar to CLIP, utilizing pre-trained pLM *f*_*P*_ and DNA language model *f*_*D*_ (Fig.1) embeddings. Given *N* pairs of binding pairs *{*(*o*_*i*_, *a*_*i*_) *}*, each modality is first processed by its respective language model to extract embeddings. These embeddings are then projected into a joint latent space. The model considers *N* (*N* − 1) negative pairs within the batch and is optimized using the infoNCE loss:

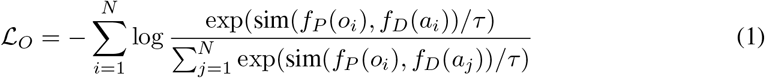

Here, sim(·, ·) is a function that computes the dot product of normalized embeddings, and *τ* is a learnable temperature parameter that scales the logits. During training, a symmetrical infoNCE loss is applied to both protein and DNA modalities, making the total training loss *L* = (ℒ _*O*_ + ℒ _*A*_)*/*2, where ℒ _*A*_ is the analogous loss calculated for the DNA embeddings. Post training, DPAC can provide a score *f* (·, ·) given any pair of protein and DNA sequence, thus enable comparing, ranking, and virtual screening.

### Simulated Annealing

Classic sampling algorithms, such as simulated annealing (12), can integrate our model to guide the sampling for binder sequence generation, as illustrated in Fig. 1. Simulated annealing is a stochastic method used to solve complex optimization problems by mimicking the physical heating and annealing process. Formally, at each sampling iteration, candidate *j* is either accepted or rejected based on the corrected Metropolis condition:

**Figure 1.**
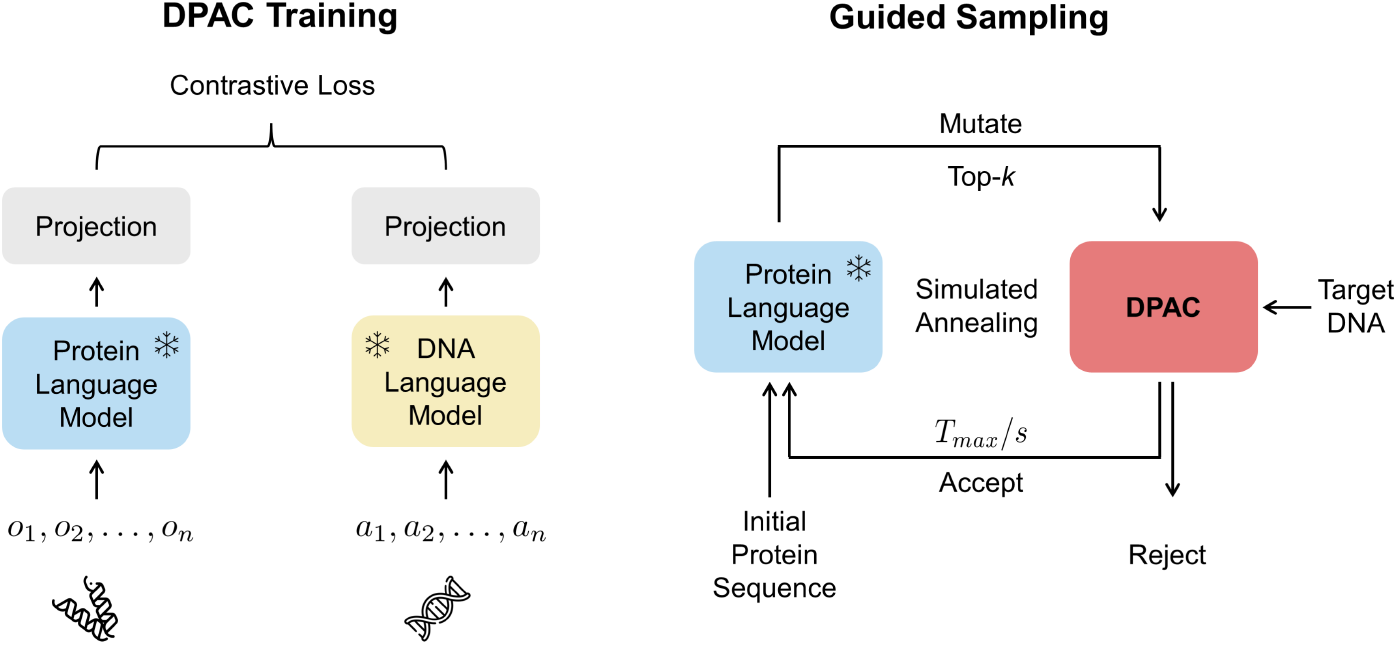
Overview of the DPAC Training and Guided Sampling Procedures. (Left) DPAC is a dual-tower model utilizing a Protein Language Model and a DNA Language Model. These models extract embeddings from input sequences, which are then projected into a joint latent space to compute a contrastive loss. (Right) Integration of the Protein Language Model in a simulated annealing framework to guide the mutation and selection of protein sequences for target DNA binding. The process iterates until a protein sequence meeting specific criteria is identified, leveraging the DPAC score for sequence evaluation.

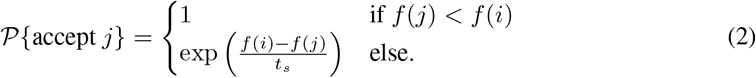

Here, *f* (·) represents the energy of the current state, and *t*_*s*_ is the temperature at step *s*. Hie et al. demonstrated that combining simulated annealing with pLM embeddings can generate valid protein sequences with hierarchical structure constraints (13). In our approach, we replace the energy optimization term with the DPAC score, defined as 1− *s*(*o*_*i*_, *a*_*i*_), for a given pair of protein and DNA sequences. We employ fast simulated annealing (14), where *t*_*s*_ is simply *T*_max_*/s*, with *T*_max_ being a hyperparameter divided by step *s*. As outlined in 1, each iteration involves randomly mutating one amino acid with ESM-2 using top-k sampling. Rather than randomly mutating across 20 amino acids, this method sets implicit constraints to produce naturalistic protein sequences while still allowing for controlled randomness in mutations. The mutated sequence is then accepted or rejected based on the criterion. The sampling loop can terminate early if (1) the score stops improving for a certain number of iterations, or (2) the best sequence reaches the mutation threshold.

#### Algorithm 1

Simulated Annealing for Protein-DNA Binding

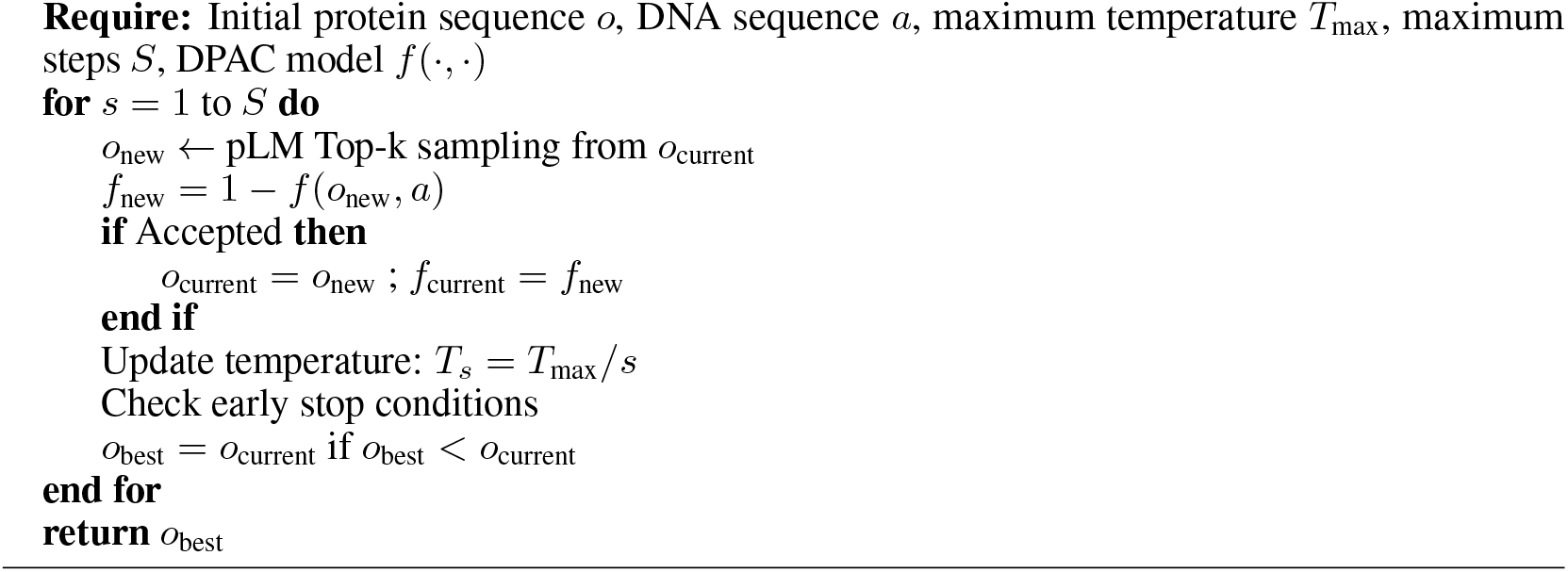

### Data

For training and evaluation, we collected DNA-protein binding data from BioLip2 (15), a semi-manually curated database with high-quality ligand-protein binding interactions. The initial dataset contains 23,817 entries. After removing duplicates, DNA motifs shorter than 3-mers, and protein sequences longer than 500 amino acids, we performed clustering for protein and DNA separately using MMSeq2 (16) with an identity threshold of 0.7. This process resulted in a strictly defined low identity set where proteins and DNAs interact only within their respective clusters. The set comprises 99 pairs of complexes, including 54 unique DNA motifs and 58 protein sequences, contributing to a library of 9,801 entries (3,132 non-redundant library). Additionally, we randomly sampled 1,000 pairs from the remaining data to create a high identity set for evaluation, bringing the total size of our training dataset to 11,419 entries.

## 3 Experiments

In this section, we explore how various protein and DNA encoding and embedding strategies influence model performance within our framework. We also detail our optimal model configuration through temperature analysis, demonstrating that our performance remains stable across changes in sequence identity. Additionally, we benchmark our model against real binding affinity measurements *K*_*D*_, observing a significant correlation. In our sampling experiments, we optimize sequences at different mutation rates to enhance the DPAC score. A case study on zinc finger nucleases illustrates that our approach can restore approximately 20% of the binding affinity *in silico*, underscoring the potential of our model in practical applications.

### 3.1 DPAC Model Design and Performance

#### Encoding Strategy

For the protein encoder, we selected ESM-2-650M, the most well-validated pLM experimentally (17; 18; 19; 20; 21; 22; 23; 24), and for the DNA encoder, we evaluated the performance of Nucleotide Transformers (NT) version 2 (500M Multi-species) (9) and DNABERT-2 (DB) (25), both trained on multi-species datasets to potentially enhance generalization. Despite freezing the encoder models, the utilization of pre-trained embeddings significantly influences tasks such as retrieval and ranking. The strategies compared include taking the maximum, average, CLS tokens, or first to last layer average (fl) (26; 27). Optimizing the combination of embeddings can significantly bridge the modality gap and establish a more uniform joint space.

We assessed model performance using AUC scores (under Receiver Operating Characteristic curve), Top 10 hit rates (HR@10), and enrichment factors at 5 and 10% (EF@0.05 and EF@0.1). For proteins, the AUC or hit rate indicates the ability to retrieve matching DNA motifs given a protein, and similarly for DNA motifs. As detailed in Table 1, in the low identity set, NT consistently outperformed DNABERT-2 across different types of embeddings. In the high identity set, various combinations of NT embeddings showed similar performance (Appendix Table 5). To enhance performance on potentially out-of-distribution samples, particularly those with low sequence identity, we opted for a combination of maximal values from ESM-2 embeddings and average values from NT embeddings.

#### Temperature Analysis

The temperature parameter *τ*, which modulates the range of logits, provides a hardness-aware property in contrastive learning (Eq. 1). Essentially, *τ* governs the mining of hard negative samples and the uniformity of the joint latent space. Typically, a smaller *τ* value intensifies the focus on mining harder negative samples, potentially enhancing representation learning. However, excessively low values can disrupt the semantic uniformity of the joint latent space (28). In our experiments, we evaluated a range of *τ* values to find the optimal setting for our prior encoding strategies, specifically ESM-max and NT-ave. Previous studies typically employed a *τ* of 0.07, as it is commonly used in CLIP-like models (11). In our tests, we explored *τ* values from 0.01 to 0.3. We observed that smaller values generally resulted in slower convergence, while larger values led to quicker convergence to a higher training losses (Figure 2a^2^). The values 0.07 and 0.09 demonstrated superior performance, prompting further investigation within this range. Ultimately, a *τ* of 0.085 yielded the best performance in our tests, as detailed in Table 2.

#### Baseline

RFAA (3) and RFNA (29) were chosen as baselines because they can co-fold proteins with DNA or RNA, making them among the most capable models available ^3^. We benchmarked both methods on our low-identity set, using a non-redundant library and the mean pairwise aligned error (PAE) to score virtual-screening performance. Our model produced substantially higher enrichment factors than RFAA and RFNA while matching their accuracy on all other metrics (Figure 2b; Tables 3 and 5).

**Table 1:** Performance comparison of different encoding strategies on Low Identity Set.

| Protein | DNA | Prot AUC | Prot HR@10 | EF@0.1 | EF@0.05 | NA AUC | NA HR@10 |
| --- | --- | --- | --- | --- | --- | --- | --- |
| ave | NT-ave | 0.521 | 0.273 | 1.028 | 1.344 | 0.528 | 0.111 |
| max | NT-ave | **0.583** | **0.404** | 1.779 | **1.581** | **0.591** | 0.141 |
| cls | NT-ave | 0.522 | 0.354 | **2.293** | 3.083 | 0.516 | 0.121 |
| fl | NT-ave | 0.53 | 0.293 | 1.146 | 0.791 | 0.538 | 0.121 |
| ave | NT-max | 0.534 | 0.303 | 1.463 | 1.107 | 0.525 | 0.081 |
| cls | NT-max | 0.529 | 0.364 | 0.83 | 1.028 | 0.535 | 0.172 |
| max | NT-max | 0.525 | 0.313 | 1.186 | 1.107 | 0.504 | 0.141 |
| fl | NT-max | 0.517 | 0.333 | 1.344 | 1.186 | 0.507 | 0.081 |
| ave | NT-cls | 0.448 | 0.111 | 0.751 | 0.158 | 0.435 | 0.09 |
| max | NT-cls | 0.515 | **0.404** | 1.304 | 1.107 | 0.516 | 0.131 |
| cls | NT-cls | 0.532 | 0.242 | 0.87 | 0.949 | 0.528 | 0.141 |
| fl | NT-cls | 0.474 | 0.212 | 1.264 | 0.158 | 0.441 | 0.101 |
| ave | NT-fl | 0.448 | 0.242 | 1.818 | 1.502 | 0.461 | 0.081 |
| max | NT-fl | 0.531 | 0.374 | 1.897 | 2.293 | 0.526 | 0.162 |
| cls | NT-fl | 0.447 | 0.282 | 1.067 | 0.949 | 0.454 | 0.111 |
| fl | NT-fl | 0.431 | 0.283 | 1.463 | 1.739 | 0.445 | 0.051 |
| ave | DB-ave | 0.509 | 0.212 | 0.751 | 0.158 | 0.479 | 0.152 |
| max | DB-ave | 0.419 | 0.232 | 0.791 | 0.712 | 0.41 | 0.081 |
| cls | DB-ave | 0.565 | 0.313 | 1.106 | 0.632 | 0.529 | 0.141 |
| fl | DB-ave | 0.488 | 0.182 | 0.395 | 0.158 | 0.462 | 0.121 |
| ave | DB-max | 0.457 | 0.242 | 0.593 | 1.107 | 0.455 | 0.141 |
| cls | DB-max | 0.516 | 0.333 | 1.699 | 1.027 | 0.525 | 0.161 |
| max | DB-max | 0.436 | 0.242 | 0.83 | 0.948 | 0.456 | 0.051 |
| fl | DB-max | 0.475 | 0.262 | 0.632 | 0.395 | 0.455 | 0.121 |
| ave | DB-cls | 0.501 | 0.404 | 1.502 | 1.344 | 0.492 | 0.171 |
| max | DB-cls | 0.453 | 0.282 | 0.988 | 0.711 | 0.455 | 0.111 |
| cls | DB-cls | 0.563 | 0.273 | 0.869 | 0.553 | 0.544 | 0.131 |
| fl | DB-cls | 0.514 | 0.353 | 1.502 | 1.66 | 0.496 | **0.191** |
| ave | DB-fl | 0.485 | 0.242 | 0.672 | 0.395 | 0.465 | 0.151 |
| max | DB-fl | 0.403 | 0.262 | 0.988 | 0.158 | 0.398 | 0.06 |
| cls | DB-fl | 0.568 | 0.353 | 1.264 | 0.948 | 0.545 | 0.151 |
| fl | DB-fl | 0.477 | 0.272 | 0.909 | 0.869 | 0.451 | 0.121 |

**Table 2:** Performance comparison of initial Temperature on Low Identity Set.

| Initial T | Prot AUC | Prot HR@10 | EF@0.1 | EF@0.05 | NA AUC | NA HR@10 |
| --- | --- | --- | --- | --- | --- | --- |
| 0.01 | 0.433 | 0.222 | 0.593 | 0.553 | 0.415 | 0.081 |
| 0.03 | 0.439 | 0.282 | 1.225 | 1.502 | 0.417 | 0.111 |
| 0.05 | 0.553 | 0.343 | 1.423 | 2.293 | 0.559 | 0.111 |
| 0.07 | 0.583 | 0.404 | 1.779 | 1.581 | 0.591 | 0.141 |
| 0.085 | 0.617 | **0.414** | 1.699 | **1.976** | **0.607** | **0.232** |
| 0.09 | **0.622** | 0.383 | 1.7 | 1.581 | **0.603** | 0.222 |
| 0.1 | 0.61 | 0.424 | 1.66 | 1.739 | 0.6 | 0.212 |
| 0.2 | 0.541 | 0.364 | 1.7 | 1.423 | 0.545 | 0.162 |
| 0.3 | 0.516 | 0.222 | 1.344 | 1.502 | 0.516 | 0.131 |

**Table 3:** Comparison of DPAC, RFAA and RFNA on Low Identity Set.

| Model | Prot AUC | Prot HR@10 | EF@0.1 | EF@0.05 | NA AUC | NA HR@10 |
| --- | --- | --- | --- | --- | --- | --- |
| RFAA | 0.546 | 0.276 | 1.111 | 0.811 | 0.541 | 0.204 |
| RFNA | 0.456 | 0.121 | 0.606 | 0.608 | 0.424 | 0.204 |
| DPAC | 0.617 | 0.414 | 1.699 | 1.976 | 0.607 | 0.232 |

#### Robustness to Sequence Identity Shift

For models leveraging sequence language models, verifying performance across sequence identities is crucial. We delved deeper into this aspect by examining changes in the AUC score relative to both protein and DNA sequence identities. Unlike the initial clustering, here each sequence’s identity was calculated pairwise against all sequences in the training set using Biotite (32).^4^ As depicted in the Fig.2c, the AUC scores are plotted against protein identity, DNA identity, and the maximum identity among the pairs of protein and DNA sequences. Notably, we did not observe a consistent pattern between AUC scores and sequence identity; performance varied across different identity levels, exhibiting both highs and lows. This variability suggests that our model maintains a certain level of robustness across sequence identity, which is beneficial for scoring interactions between unseen proteins and DNA.

### 3.2 Binding Affinity Benchmarking with DPAC Scores

We evaluated binding–affinity prediction on the protein–nucleic-acid subset of PDBBind v2020 (33), excluding all NMR structures for consistency. Because absolute *K*_*D*_ values are highly sensitive to assay conditions, we compared only measurements obtained under identical experimental setups. We treated each mutant-series experiment, where a single protein is tested against multiple DNA motifs as an independent evaluation group. The resulting benchmark comprises 68 groups. Each group contains one protein sequence paired with 2 to 17 DNA motifs (28 groups have more than two motifs). For every protein–motif pair we computed a DPAC logit score and, within each group, fitted a simple linear regression of logit score versus reported affinity (nM). A negative slope indicates correct ranking (higher logit → lower *K*_*D*_). DPAC achieved negative slopes in 55 of 68 groups (80.8%). A permutation test over group assignments yielded *p* = 9 × 10^*−*5^, providing strong evidence against the null hypothesis of no association. Figure 2d illustrates eight representative cases, four with negative and four with positive slopes, highlighting the range of relationships observed. Overall, the predominance of negative correlations demonstrates that DPAC logits are a reliable proxy for protein–DNA binding strength.

### 3.3 Simulated Annealing with DPAC

We applied simulated annealing guided by DPAC logits to improve binding scores while keeping sequence changes within realistic limits. The protocol used top-3 sampling, a maximum temperature of 50, and up to 500 steps. Three mutation caps were tested (10%, 30%, and 50%) of positions and each optimisation was repeated three times per sequence in the low identity set. To gauge naturalness of resulting proteins, we estimated sequence likelihood with ProGen2 (34) perplexity. As the baseline we followed the recent DNA-binder study (35) and ran LigandMPNN (36) on the same RFNA/RFAA-folded complexes, redesigning each protein at temperatures 0.1, 0.2, and 0.3 (three sequences per target). For parity we generated the same number of DPAC-optimised sequences by running the simulated-annealing pipeline three times. Both sets were scored with DPAC logits (binding) and ProGen2 perplexity (naturalness). DPAC-guided optimisation boosted binding predictions to a level comparable with LigandMPNN, particularly at the 10% mutation limit, while both methods showed higher perplexity than the wild-type sequences. These results reinforce DPAC’s effectiveness as an optimization objective alongside a state-of-the-art binder-design approach.

For a practical application, we selected a classic zinc finger protein, Zif268 (PDB ID 1AAY) (37), commonly used in previous zinc finger nuclease (ZFN) design studies (38), and thus representing a key genome editing scaffold (39). Instead of starting with a binder and optimizing it, we first introduced random amino acid mutations at rates of 10%, 30%, and 50% to corrupt the original sequence. We then assessed how much binding affinity our optimization approach could restore. The *in silico* binding affinity was measured using the AlphaFold 3 server, the latest model that incorporates a diffusion model for all-atom folding, thus including DNA-protein complexes (4). Specifically, we used the predicted template modeling (pTM) score for overall complex structure confidence and the interface predicted template modeling (ipTM) score for binding interface confidence. The results are displayed in Table 4 and Figure 3. Red represents mutated sites, and yellow in the optimized proteins indicates recovered amino acids that match those in 1AAY. It is important to note that red in the corrupted and optimized proteins does not necessarily indicate the same amino acids (see Appendix Section G for exact sequence details). For the 10% corruption, since the mutated region is small and not pivotal for binding from a structural perspective, both the optimized and corrupted sequences exhibit similar ipTM scores to 1AAY. However, for 30% and 50% corruption, which disrupt the interaction between the third alpha-helix and the DNA minor groove, our approach showed promising recovery of binding affinity around 20%. After sampling, binding was reconstructed with optimized amino acids.

**Figure 2.**
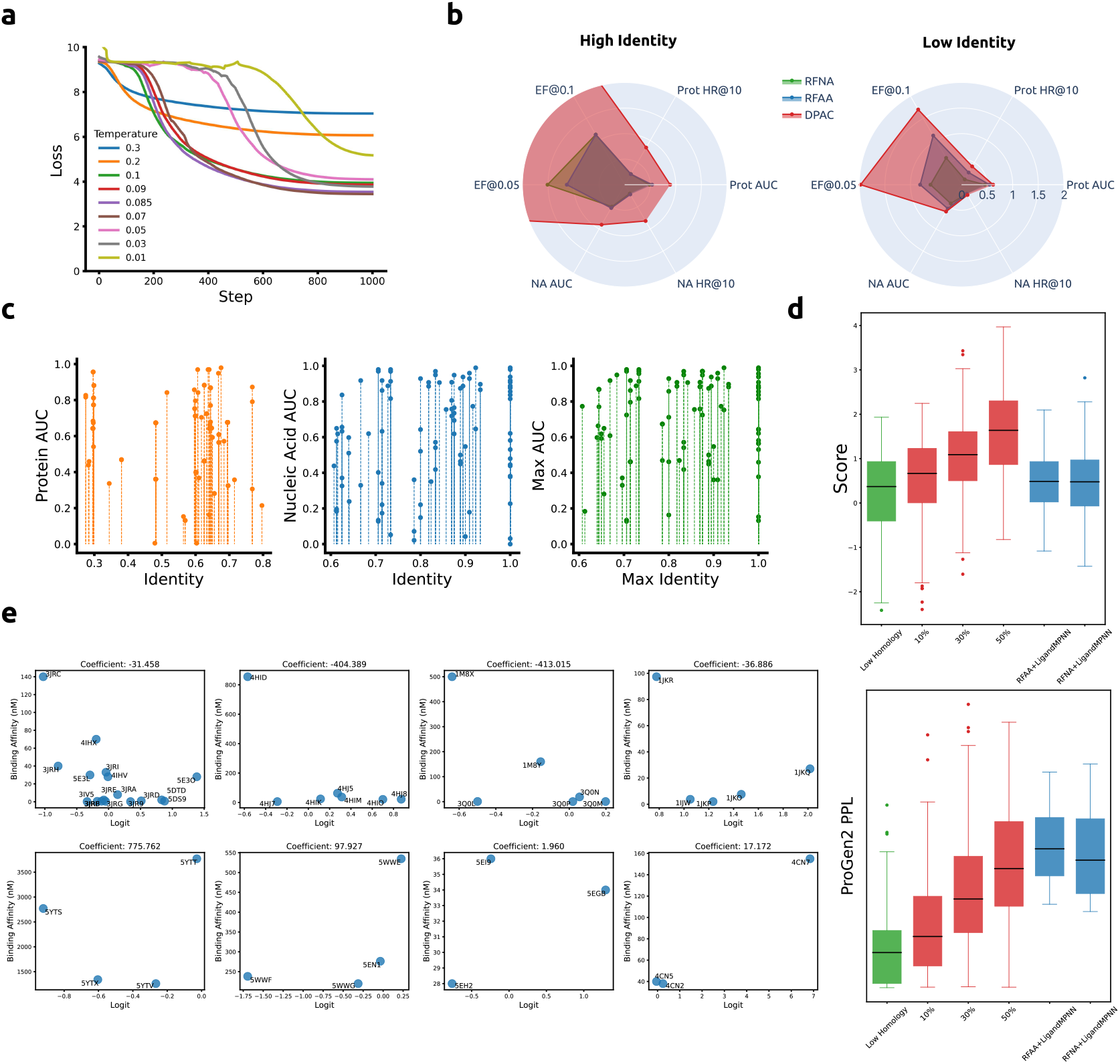
(a) Training loss and initial Temperature. (b) Comparison of DPAC, RFAA and RFNA on different metrics. (c) Robustness analysis across sequence identities. AUC scores against sequence identity for proteins (left, orange), nucleic acids (center, blue), and the maximum identity of protein-DNA pairs (right, green). (d) Examples of Binding Affinity Regression. Four examples each of positive (Top) and negative (Bottom) correlations between DPAC Logit scores and Binding Affinity (nM). Pearson correlation coefficients are displayed at the top of each plot. (e) Optimization Benchmark. Protein sequence optimization was performed on the low identity set (green). Each set was evaluated using DPAC logits as binding scores and ProGen2 perplexity. Our DPAC-optimized groups (blue) were generated with different mutation constraints (10%, 30%, 50%). Two additional groups (red) utilized LigandMPNN to optimize DNA-protein complexes folded by either RFAA or RFNA.

**Table 4:** 1AAY binding recovery.

| Corruption | Optimized |  |  | Corrupted |  |  |
| --- | --- | --- | --- | --- | --- | --- |
|  | DPAC Score | ipTM | pTM | DPAC Score | ipTM | pTM |
| 10% | 0.817 | 0.91 | 0.87 | 0.381 | 0.91 | 0.86 |
| 30% | 0.824 | 0.89 | 0.84 | 0.459 | 0.77 | 0.54 |
| 50% | 1.641 | 0.83 | 0.62 | 0.559 | 0.66 | 0.4 |
| 1AAY (0%) | - | - | - | 0.8913 | 0.93 | 0.9 |

**Figure 3.**
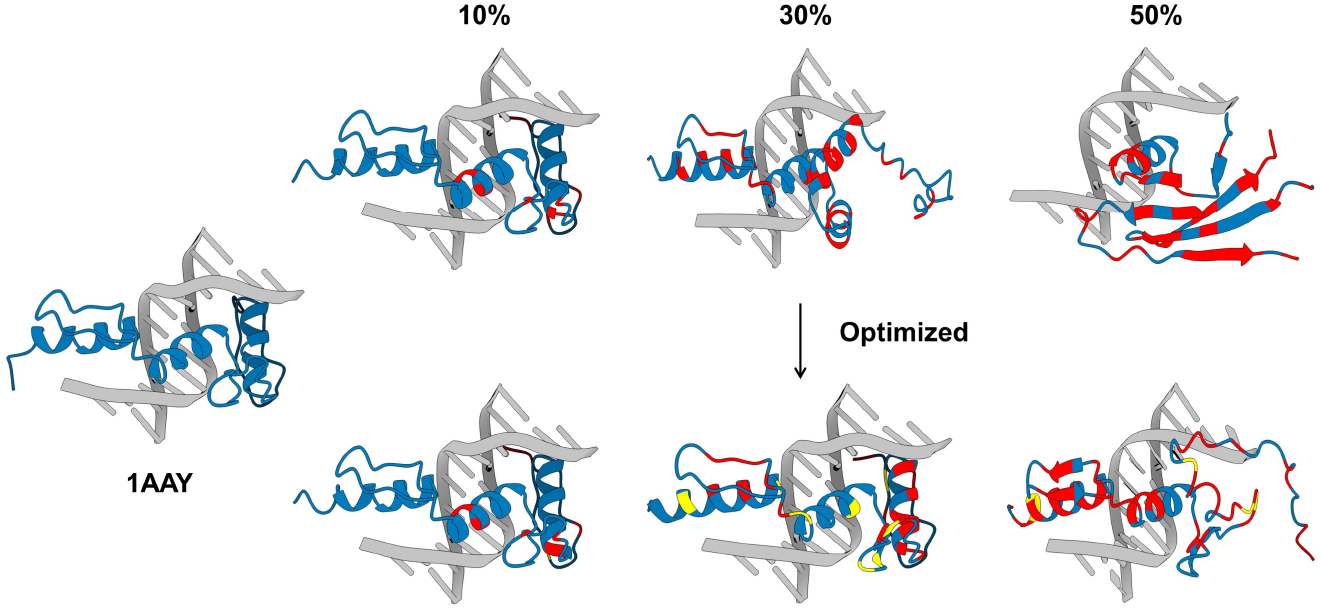
Visualization of mutation and optimization effects on ZFN (1AAY). The first column shows the original structure, followed by subsequent columns representing increasing mutation rates. Blue indicates the unchanged regions of the protein, red highlights areas with mutations, and yellow marks the recovery of original amino acids in the optimized structure. The bottom row demonstrates the final optimized protein structures across different mutation rates, showcasing the capability of our optimization method to restore binding affinity even after significant sequence disruption. All structures were folded on the AlphaFold 3 server.

## 4 Conclusion

In this work, we introduce DPAC, a novel sequence-based contrastive learning framework for predicting and designing protein-DNA interactions. By aligning pre-trained protein and DNA language models in a shared latent space, DPAC enables efficient and scalable modeling of binding interactions, outperforming state-of-the-art structure-based methods on a low identity dataset. Moreover, DPAC integrates simulated annealing to guide the optimization of protein sequences for enhanced DNA binding. A case study on zinc fingers demonstrates the effectiveness of this approach, recovering up to 20% of binding affinity *in silico* after significant sequence disruption. DPAC represents an important step towards purely sequence-based modeling and design of protein-DNA interactions, offering a more computationally efficient alternative to existing structure-based methods. However, there are still limitations and areas for future improvement, including enhancing performance on low identity sequences, incorporating post-translational modifications (PTMs) for enhanced DNA binding (40), and exploring alternative generative methods for sequence design, for example guided discrete diffusion (41; 42; 43) and discrete flow matching models (44; 45). Nonetheless, our current DPAC model provides a powerful new tool for studying and engineering protein-DNA interactions at the sequence level, unlocking new possibilities for wide-scale genomic applications, including precision genome editing and targeted gene regulation.

## 5 Declarations

## Acknowledgments

We thank the Duke Compute Cluster, Pratt School of Engineering IT department, and Mark III Systems, for providing database and hardware support that has contributed to the research reported within this manuscript.

## Author Contributions

L.T.C. devised model architectures and theoretical formulations, and trained and benchmarked models. R.P. and P.V. assisted in data collection and CLIP model training. P.C. designed, supervised, and directed the study, and finalized the manuscript.

## Data and Materials Availability

Model and inference code can be found at https://github.com/programmablebio/dpac.

## A Related Works

### Contrastive Learning and Biology Applications

Contrastive learning has been widely used to align different modalities in a joint latent space, especially texts and images, and shows successful and robust performance across various retrieval tasks (11; 46; 47). The contrastive-based approach has recently introduced into biological application for different molecular modalities, including peptides, protein, DNA, and small molecules. As an example, ConPLex predicts drug-target interactions via triplet distance loss, focusing specifically on distinguishing true drug-target pairs from decoy compounds (48). DrugCLIP (49) and PepPrCLIP (18) leverage a CLIP-like structure and infoNCE loss for small molecule and peptide binder retrieval tasks given a target protein. Finally, CLAPE, a protein-DNA binding site prediction model, uses triplet center loss with pLM to identify DNA binding residues (50).

### Protein DNA Interaction Modeling

Computational modeling of protein-DNA interactions can be broadly categorized into two main approaches: contact prediction and structure folding. In the contact prediction approach, the objective is to predict whether amino acids on an input protein sequence are DNA binding sites or not via a binary classification task. Classical methods typically incorporate both protein sequence and structure information using graph neural networks (GNNs). For instance, GraphSite (51) employs a graph transformer architecture to model contact prediction as a node classification problem. Similarly, GraphBind (52) constructs hierarchical graph neural networks to embed local patterns for improved prediction accuracy. More recently, EquiPNAS (53) leverages protein language models in conjunction with equivariance graph neural networks to enhance binding prediction performance. On the other hand, the folding approach aims to directly model the three-dimensional structure of protein-nucleic acid complexes. Recent efforts from the RosettaTTAFold and AlphaFold teams have extended their previous protein structure prediction models to accommodate the folding of protein-nucleic acid complexes. These advancements have led to the development of RFNA(29), RFAA (3), and AlphaFold 3(4), which have shown promising results in predicting the structures of protein-DNA complexes.

### *De Novo* Binder Design

Generative protein binder design with new deep learning techniques fall primarily into either strucure-based pipelines or sequence-based language modeling methods. RFDiffusion generates binders to target proteins by employing a diffusion-based method (54), to iteratively refine initial protein structures, optimizing their conformations and interactions for high-affinity binding to the specified target proteins (55). RFAA leverages the same generative procedure to design protein binders to nucleic acids and small molecule compounds (3). EvoPlay (56) uses self-play reinforcement learning with AlphaFold2 (57) for peptide binder design. For peptide binder design directly from target sequence, the recent PepMLM model leverages span masked language modeling to mask and unmask new binders at the C-terminus of target protein sequences (19). Finally, recent studies have demonstrated the possibility of designing DNA motif-specific binders using classical methods as the Rotamer Interaction Field (RIF) docking and generation (35).

### Protein Sequence Sampling with pLM

With the rise of pLMs, recent efforts have shown that protein sequences can be directly sampled from pLMs. Plug and Play Directed Evolution (58) leverages gradient-based discrete Markov Chain Monte Carlo (MCMC) to sample mutations from the joint distribution of a supervised classifier and general pLMs. For gradient-free methods, it has been demonstrated that the combination of pLMs and simulated annealing can generate valid protein sequences under hierarchical structural constraints (13). While this approach offers greater flexibility, it is more computationally expensive compared to gradient-based methods (59).

## B High Homology Set Performance

In this section, we examine the performance of various encoding strategies (Table 5) and the impact of different temperature settings (Table 6) on the high identity set. Regarding the combinations of encoding methods, most pairings between ESM and NT consistently deliver strong results, achieving AUC scores above 80% and hit rates around 60%. However, DNABERT-2 (10) exhibits less impressive performance across its various encoding types. In our temperature studies, we found that both extremes (too low (0.01) and too high (0.3)) as anticipated, disrupted the uniformity of the joint latent space, leading to poorer performance compared to other temperature settings.

**Table 5:**
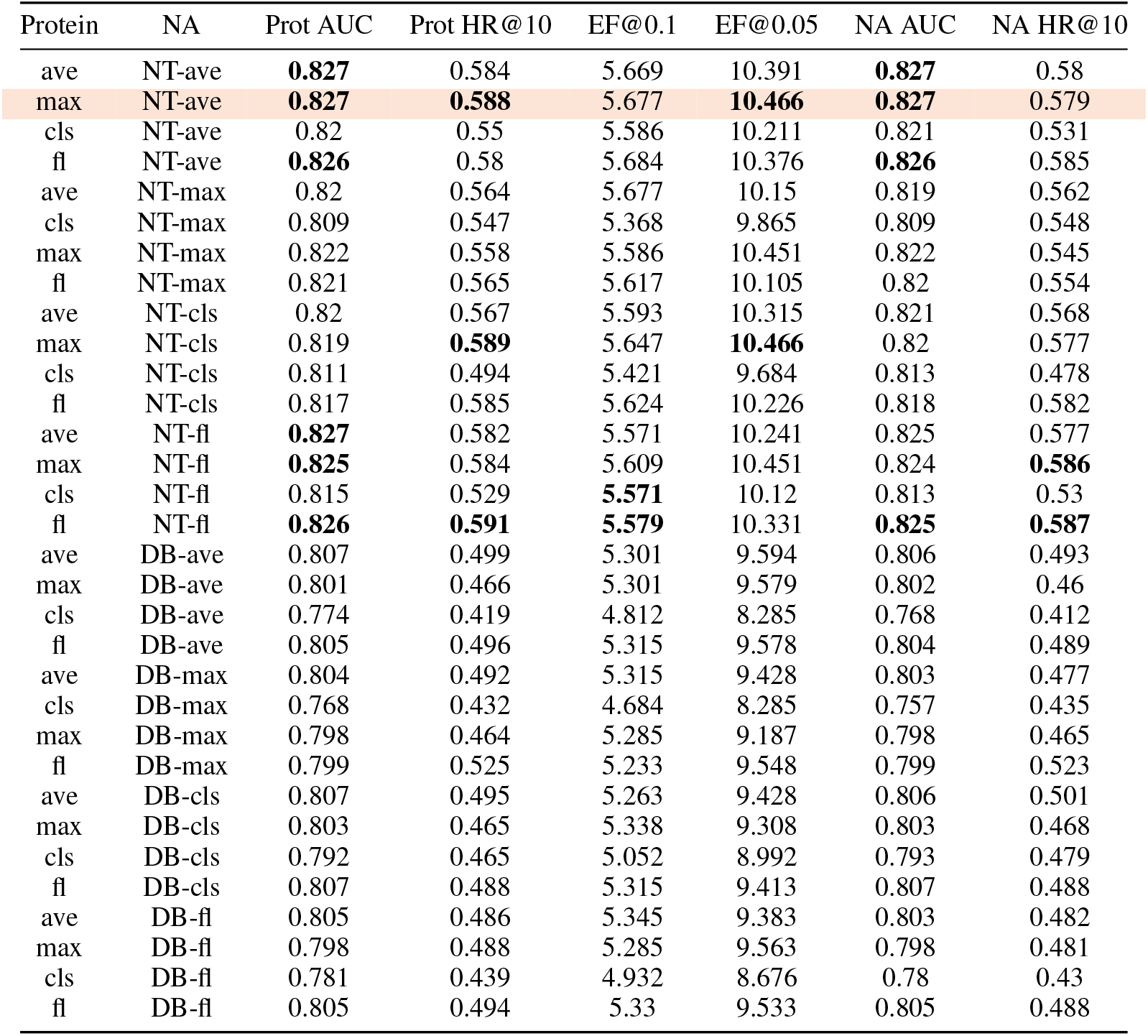
Performance comparison of different encoding strategies on High Homology Set.

**Table 6:**
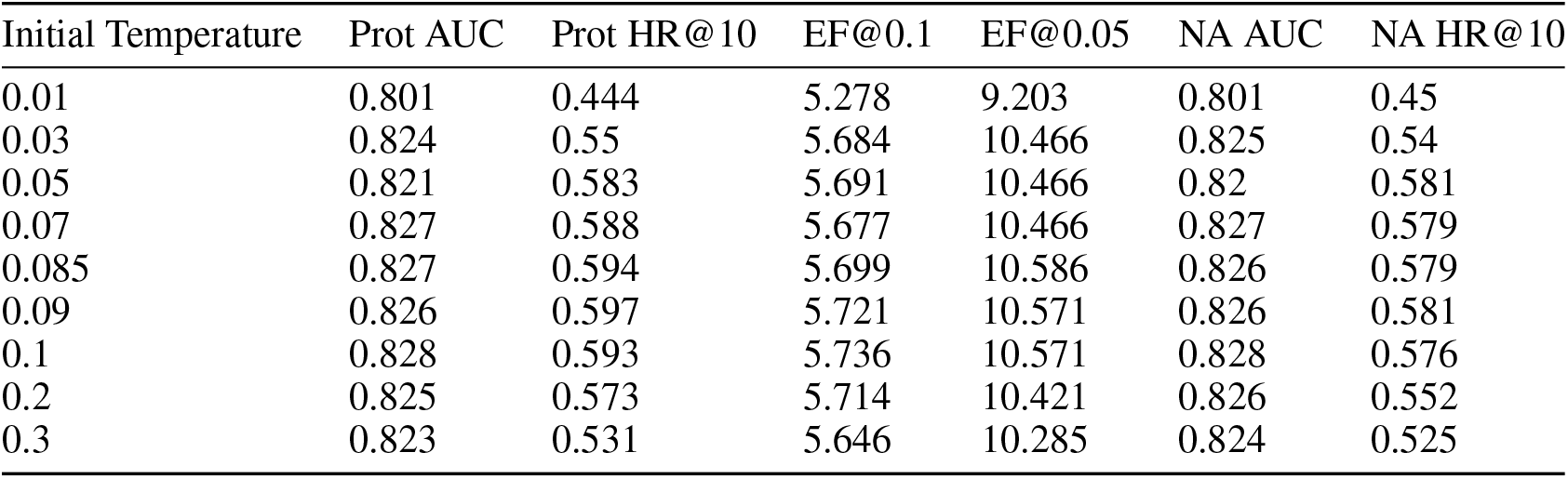
Performance comparison of Initial Temperature on High Homology Set.

| Initial Temperature | Prot AUC | Prot HR@10 | EF@0.1 | EF@0.05 | NA AUC | NA HR@10 |
| --- | --- | --- | --- | --- | --- | --- |
| 0.01 | 0.801 | 0.444 | 5.278 | 9.203 | 0.801 | 0.45 |
| 0.03 | 0.824 | 0.55 | 5.684 | 10.466 | 0.825 | 0.54 |
| 0.05 | 0.821 | 0.583 | 5.691 | 10.466 | 0.82 | 0.581 |
| 0.07 | 0.827 | 0.588 | 5.677 | 10.466 | 0.827 | 0.579 |
| 0.085 | 0.827 | 0.594 | 5.699 | 10.586 | 0.826 | 0.579 |
| 0.09 | 0.826 | 0.597 | 5.721 | 10.571 | 0.826 | 0.581 |
| 0.1 | 0.828 | 0.593 | 5.736 | 10.571 | 0.828 | 0.576 |
| 0.2 | 0.825 | 0.573 | 5.714 | 10.421 | 0.826 | 0.552 |
| 0.3 | 0.823 | 0.531 | 5.646 | 10.285 | 0.824 | 0.525 |

## C Implementation Details

Our final model employs two separate multi-layer perceptrons (MLPs) for processing protein and nucleic acid embeddings. Each branch consists of a linear projection layer, followed by GELU activation and a subsequent projection that reduces it to the target dimension. Layer normalization is applied after the final projection to enhance training stability. The embeddings for proteins and nucleic acids are sourced from pre-trained models ESM2-650M and NucleotideTransformer2-500M, respectively, with dimensions of 1280 for proteins and 1024 for nucleic acids (7; 9). The joint latent space has dimension of protein embeddings 1280. The temperature was set to be 0.085. The model is trained over 1000 epochs using the AdamW (60) optimizer with an initial learning rate of 1 × 10^− 4^ and a batch size of 11,418 on single NVIDIA GPU DGX A100-80G. A cosine annealing schedule is employed to adjust the learning rate across epochs with minimum learning of 1 × 10^− 6^.

## D Evaluation Metrics

### Hit Rate

The hit rate is a measure used to evaluate the performance of a predictive model in ranking. Specifically, the top-10 hit rate indicates the proportion of times the correct binder or ligand is identified within the top 10 predictions of the model.

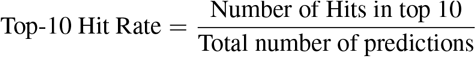

where a “Hit” is counted when the actual relevant binder or ligand is found within the top 10 predictions.

### Enrichment Factor

The enrichment factor (EF) is a metric used commonly to assess the ability of a method to enrich relevant binders or ligands within a subset of its predictions. It is particularly useful in scenarios like virtual screening. The EF at a certain percentage (e.g., 5% or 10%) compares the concentration of relevant molecules in the top-ranked subset of predictions to their concentration in the overall dataset. For a given percentage *x*, the enrichment factor *EF*_*x*_ can be expressed as:

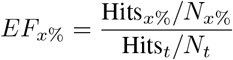

where where Hits_*x*_% represents the number of true binding molecules (hits) in the top *x*% of the predicted molecules, *N*_*x*_% is the total number of molecules in the top *x*%, Hits_*t*_ denotes the total number of true binding molecules in the entire dataset, and *N*_*t*_ represents the total number of molecules in the dataset.

### PAE

The Predicted Aligned Error (PAE) is a metric that estimates the error in the relative position and orientation in the predicted structure. It is calculated by aligning each frame (i) and computing the errors over all C*α* (or P atom) coordinates. For atom nodes, the canonical frames are aligned, and the error is computed over all other nodes (3). Higher PAE values indicate higher errors and lower confidence in the predicted structure.

### pTM and ipTM

The predicted template modeling (pTM) and interface predicted template modeling (ipTM) scores derive from the template modeling (TM) score, which measures structural accuracy. A pTM score above 0.5 suggests the predicted fold might be similar to the true structure. The ipTM score assesses the predicted positions of subunits within a complex. Scores above 0.8 indicate high-quality predictions, while scores below 0.6 suggest likely failures. Scores between 0.6 and 0.8 are ambiguous (4).

### Perplexity

Perplexity measures how well a probabilistic model predicts a sample and is commonly used in language models, including protein language models, to evaluate sequence. The perplexity is defined as the exponential of the average negative log-likelihood per token.

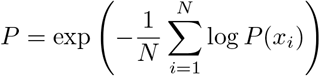

where *N* is the number of tokens, and *P* (*x*_*i*_) is the predicted probability of the *i*-th token. Lower perplexity indicates better model confidence. For protein language models like ProtGPT2, perplexity is found to be associated with structural confidence: lower perplexity suggests a higher likelihood of the model producing valid protein structures (61).

## E Efficiency Analysis

The computational efficiency of both RFAA and DPAC during virtual screening was assessed by recording the time taken for processing sample sizes of 100, 500, 1,000, and 2,000. The results are detailed in Table 7 and illustrated in Figure 4. While DPAC completes the screening of 2,000 samples within 2 minutes, RFAA requires approximately 336 hours to process the same number. Extrapolating these results to a hypothetical library of one million samples, assuming linear scaling, DPAC would complete the task in about 15 hours, whereas RFAA would need an impractical 7,000 days.

**Table 7:** Time efficiency of DPAC and RFAA.

| # of Samples | DPAC | RFAA |
| --- | --- | --- |
| 100 | 5.79 | 70,440 |
| 500 | 26.96 | 331,588 |
| 1,000 | 53.45 | 617,493 |
| 2,000 | 107.57 | 1,208,840 |

**Figure 4.**
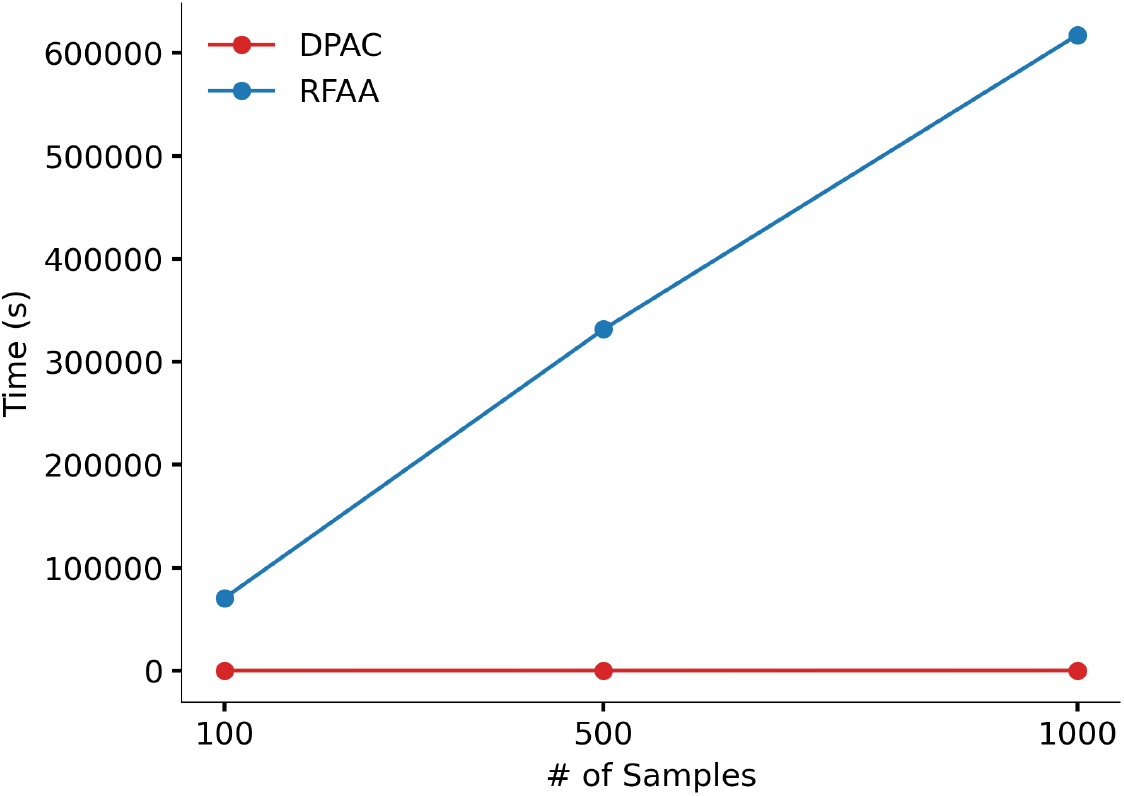
Time efficiency of DPAC and RFAA

## F Projected Dimension Study

Table 8 presents an ablation study of different projected dimensions on the Low Homology Set under the optimal configuration of ESM-max and NT-average with a temperature of 0.085. It is noteworthy that as the dimension decreases, the performance drops, particularly for AUC scores. One possible explanation for this observation is that the original dimension of ESM-2 might contain enriched features that significantly contribute to the model’s ability to generalize over sequence identities. Moreover, by maintaining the same embedding size as the protein language model (pLM), it becomes feasible to perform conditional modeling or generation by directly replacing specific token embeddings, such as the cls token, in the original protein embeddings.

**Table 8:** Comparison of different projected dimension on Low Homology set.

| Dim | Prot AUC | Prot HR@10 | EF@0.1 | EF@0.05 | NA AUC | NA HR@10 |
| --- | --- | --- | --- | --- | --- | --- |
| 128 | 0.456 | 0.404 | 1.225 | 1.028 | 0.466 | 0.061 |
| 256 | 0.513 | 0.283 | 1.146 | 1.739 | 0.496 | 0.101 |
| 512 | 0.54 | 0.363 | 1.265 | 1.265 | 0.516 | 0.121 |
| 1280 | 0.617 | 0.414 | 1.699 | 1.976 | 0.607 | 0.232 |

## G 1AAY Mutated and Sampling Sequences

Original 1AAY Sequences:

- Protein: MERPYACPVESCDRRFSRSDELTRHIRIHTGQKPFQCRICMRNFSRSDHLT-THIRTHTGEKPFACDICGRKFARSDERKRHTKIHLRQKD
- DNA1: AGCGTGGGCGT
- DNA2 (used for our sampling): TACGCCCACGC

Mutated Sequences:

- 10%: NTRPYACTVESNDRRFSRSDELTRHIRIMTGQAPFQCCICMRNFSRS-DHLTTSICTHTGEKPFACDICGRKFARSDERKRHTKIHLRQKD
- 30%: LERPKACPVESCDQRHSRSPELDRLANICKGQKPDMRRIIMKNFSRSDHIT-THIRTHQNENPKFTDICGRKFAWSDEMKRHSKIVLRQKD
- 50%: HHTPYADPQESCMRIKLRSGWLEYHIWFLPCPKPFQIRICMRNTIRS-DHLTAHVNGHFLEVFFFYDICGVFFQRSITAKRSLKIRMCCKR

Optimized Sequences

- 10%: NTRPYACTVESCDRRFSRSDELTRHIRIQTGQAPFQCCICMRNFSRSDHLTTSIC-THTGEKPFACDICGRKFARSDERKRHTKIHLRQKD
- 30%: LERPFACPVESCDSRFSRSSELDRLLNICSGQKPYQYRICMRNFSRSDHLT-THIRTHTNERPYFRDICGRKFARSDEMKRHVKIHLRQKD
- 50%: HSKPYALPQESCSRTKLRSDQLNYHIKFHPGEKPFQIRICMRNYIRSDHLTD-HVKGHSGEVFFFSDICGVSFQRSNTAKRLRKIRLRKKR

## Footnotes

1 DPAC is also the performing arts theater in Durham, North Carolina Preprint and preprint only

2 The loss curves are processed over running average window of 50 steps for better visualization.

3 The work was done before the release of AlphaFold3 model weight and other similar models (30; 31).

4 Biotite shows higher identity values due to its different pairwise alignment method as compared to MMSeqs2.

